# Differential peripheral immune dynamics underlie therapeutic response to chemotherapy and chemo-immunotherapy in triple-negative breast cancer

**DOI:** 10.64898/2026.04.28.721416

**Authors:** Zahra Mesrizadeh, Kavitha Mukund, Shankar Subramaniam

**Affiliations:** Department of Bioengineering, UC San Diego, Gilman Drive, La Jolla, CA 92093, USA; Departments of Cellular & Molecular Medicine, Computer Science & Engineering, and Data Science, UC San Diego, Gilman Drive, La Jolla, CA 92093, USA

**Keywords:** Chemotherapy, immune checkpoint blockade, triple negative breast cancer, tumor microenvironment, single-cell RNA sequencing, breast cancer, immune response

## Abstract

Triple-negative breast cancer (TNBC) remains the most aggressive breast cancer subtype, with limited treatment options and variable response to immune checkpoint inhibitors. While tumor-infiltrating lymphocytes have been extensively studied, the integration of system-level peripheral immune dynamics with mechanistic immune regulation underlying therapeutic response and resistance remain poorly defined. Here, we integrate systems-level immune state modeling with pathway-level mechanistic inference to analyze single-cell RNA sequencing of peripheral blood mononuclear cells from advanced TNBC patients treated with paclitaxel alone (chemotherapy) or in combination with anti-PD-L1 antibody atezolizumab (combination). This framework leverages treatment arm, longitudinal sampling, and clinical response to resolve coordinated immune programs across lymphoid and myeloid compartments. Using this approach, we identified distinct treatment- and response-specific immune states in pre- and post-treatment. Chemotherapy responders displayed pre-treatment adaptive immune priming, whereas combination therapy responders exhibited pre-existing effector T cell activity coupled with tumor tissue PD-L1 expression. In contrast, chemotherapy non-responders developed persistent post-treatment immune dysregulation in regulatory and terminal effector programs, while combination therapy non-responders demonstrated maladaptive remodeling of adaptive and innate lymphoid compartments, including dysfunctional NK and metabolically reprogrammed myeloid populations. Across both regimens, pathways involving protein translation, metabolic adaptation, and stress signaling emerged as critical modulators of response. These findings suggest that coordinated adaptive–innate immune dynamics underlie therapeutic efficacy, whereas systemic immune exhaustion and myeloid immunoregulation lead to resistance. Projection of these peripheral immune programs onto independent I-SPY2 showed concordant associations with tumor immune phenotypes and pathological complete response, supporting generalizability of the identified systemic immune states. Our study demonstrates the utility of an integrative systems-level approach for linking peripheral immune state organization with mechanistic insights, informing immune response and resistance in TNBC.

## 1. Introduction

Triple-negative breast cancer (TNBC) is the most aggressive subtype of breast cancer, characterized by early relapse, poor prognosis, and an absence of targeted therapy. Despite the use of standard chemotherapy, approximately one-third of patients with stage II/III TNBC develop distant metastases within 2-3 years of diagnosis (1,2). The addition of immune checkpoint inhibitors (ICIs) to chemotherapy have improved survival in several malignancies with high tumor mutational burdens (3). This prompted investigation into their efficacy in TNBC, a relatively immunogenic subtype of breast cancer, enriched for tumor-infiltrating lymphocytes (TILs). In the IMpassion130 trial, combination of atezolizumab with Nab-paclitaxel improved progression-free survival (PFS) in PD-L1 advanced TNBC patients (median PFS: 7.5 vs. 5.0 months), leading to accelerated regulatory approval (4,5). However, follow-up studies, including IMpassion131, which paired atezolizumab with paclitaxel, and a separate international phase III trial (ALEXANDRA/IMpassion030), resulted in responders and non-responders, even in PD-L1 patients (2,6). These mixed outcomes underscore a critical challenge that only a subset of TNBC patients achieve a durable response, highlighting the need to better define the system-level immune mechanisms that differentiate therapeutic response and resistance.

While most efforts to date have focused on local immune responses within the tumor microenvironment (TME), effective antitumor immunity requires continuous interplay between the TME and the peripheral immune system. Circulating immune cells reflect systemic immune competence and, importantly, offer accessible and dynamic biomarkers of treatment response and disease progression. To date, peripheral immune profiling in breast cancer has largely relied on broad metrics such as neutrophil-to-lymphocyte (NLR) or lymphocyte-to-monocyte ratios (LMR) (7,8), or limited immunophenotyping of major immune subsets using next-generation sequencing technologies such as single-cell RNA sequencing (scRNA-seq) (1). Emerging studies in TNBC suggest that systemic immune dysregulation is a defining feature. Early-stage patients exhibit reduced circulating B cells and elevated regulatory T cells compared to healthy controls (9), while patients with advanced disease demonstrate widespread immunosuppression marked by diminished CD4 T cells, expansion of monocytes, and dysfunctional effector populations (10). These studies have largely lacked integrative, systems-level analyses capable of resolving the mechanistic basis of systemic immune response and resistance, particularly after chemotherapy versus combination therapy. Large neoadjuvant trials such as I-SPY2 have generated clinically annotated transcriptomic datasets linking systemic immune signaling to treatment response but these analyses largely rely on bulk immune signatures and do not resolve high-resolution immune subset states or coordinated peripheral immune programs (11,12).

Here, we apply an integrative systems-level immune state analysis to scRNA-seq data from peripheral blood mononuclear cells (PBMC/blood) and breast tissue with advanced TNBC treated with paclitaxel alone or in combination with anti-PD-L1 antibody atezolizumab (ATZ) (combination therapy) (13). Our analysis highlights the transcriptional programs and immune cell states associated with therapeutic response (in responders and non-responders) and treatment regimens (chemo- and combination therapy). Within peripheral immune cells (blood), chemotherapy responders exhibited pre-treatment adaptive immune priming, whereas combination therapy responders showed pre-existing effector adaptive immunity with innate lymphoid activation. Chemotherapy non-responders developed persistent post-treatment immune dysregulation in regulatory and terminal effector programs, while combination therapy non-responders showed post-treatment remodeling in adaptive and innate lymphoid compartments. Together, our study delineates coordinated systemic immune programs associated with therapeutic efficacy and resistance, providing insight into peripheral regulation in TNBC.

## 2. Results

### 2.1. Identification of key mechanisms driving peripheral blood immune cell dynamics across treatment and response

Overall, 143,085 and 289,752 high-quality cells across four major cell types were identified after rigorous quality control across tissue and blood, respectively (see Methods). To systematically assess treatment-associated immune alterations, we analyzed changes in circulating immune cell frequencies post- versus pre-treatment across each treatment and response group. Given biased patient-specific clustering of immune cell types in tissue samples - except for T cells - we focused most of our dynamic analysis on blood (**Figure 1A, B**) (**Supplementary Figure S1A, B**). In chemotherapy-treated patients, responders (R) exhibited a post-treatment decrease in B cells, T cells, and myeloid cells, alongside an increase in NK cells (**Figure 1C**). In contrast, non-responders (NR) showed a global increase across all immune cell types following treatment. In the combination therapy group, both responders and non-responders displayed increased immune cell populations post-treatment, except B cells, which decreased in both groups (**Figure 1C**).

**Figure 1.**
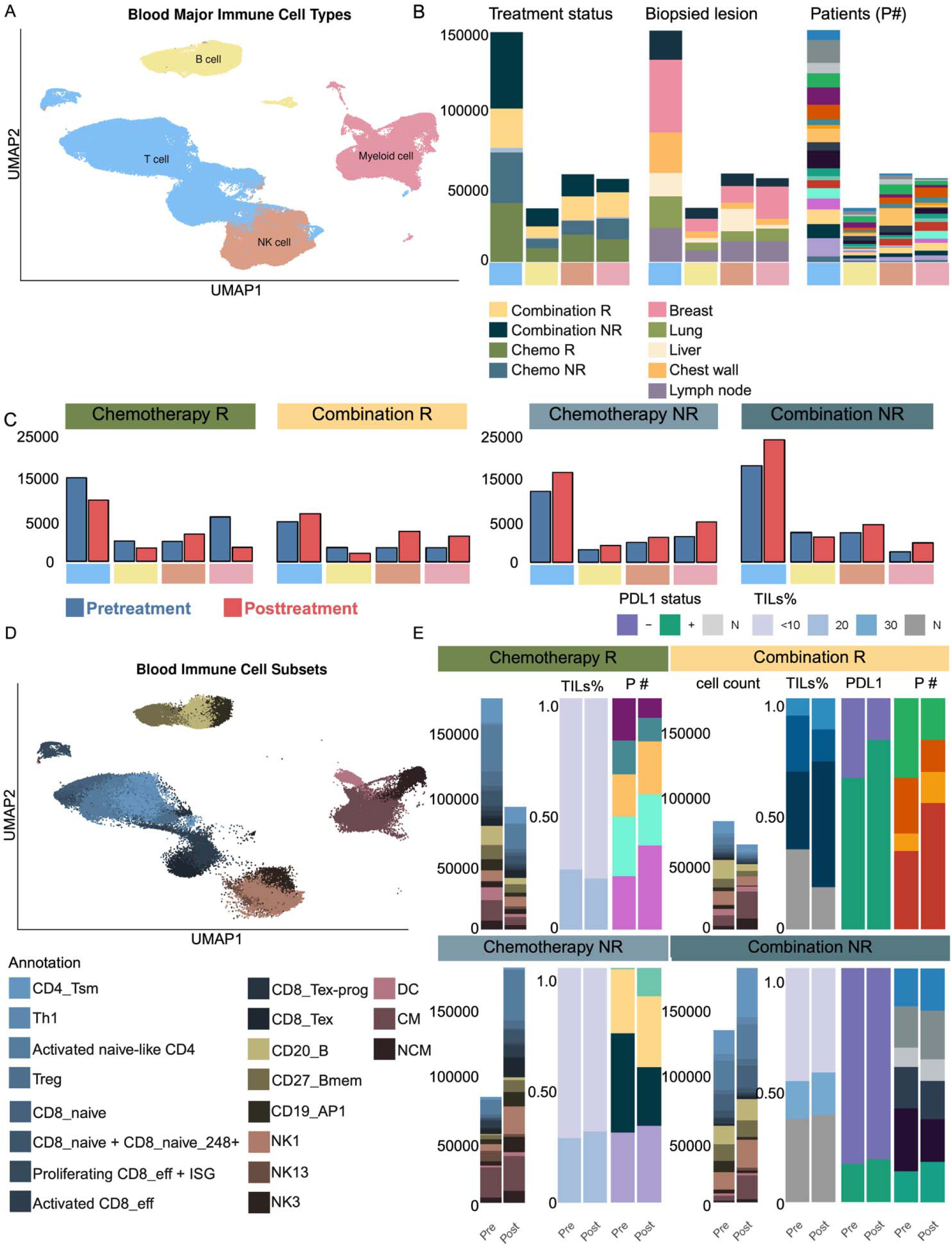
Comparative immune landscape of blood samples under different treatment conditions. **(A)** UMAP plots depict major immune cell types in blood, highlighting T cells (blue), B cells (yellow), monocytes (pink), and NK cells (brown). **(B)** Compositions of major immune cell types in treatment and response groups, biopsied lesions, and individual patients. **(C)** Bar plots quantifying immune cell frequencies, illustrating treatment-associated shifts in immune cell distribution. **(D)** UMAP plot depicting blood immune subsets, with colors denoting functionally annotated subsets. **(E**) Integrated bar plots showing frequencies of functional immune subsets, proportions of tumor-infiltrating lymphocytes (TILs), PD-L1 (PDL) cells, and patient group IDs (P#), stratified by responder (R) and non-responder (NR) status. This panel links peripheral immune subset dynamics with intratumoral immune contexture.

To resolve immune programs associated with these frequency shifts, we next stratified major immune lineages into functional sub-clusters based on gene expression programs (e.g., activation, exhaustion, migration) using known immunological signatures (**Figure 1D**) (1,2). We focused on subtypes that were (1) representative across patients and (2) differentially enriched by treatment response. All comparisons were made between post- and pre-treatment samples within each group. Although stromal TIL levels and peripheral immune cell counts often only moderately correlate in TNBC patients (3), declining TILs following chemotherapy have been linked to treatment response, while persistently high post-treatment TILs can lead to poor outcomes (4). Furthermore, the predictive value of TILs and PD-L1 expression in informing ICI efficacy is well-established (5). In this dataset, analysis of response-specific TIL levels demonstrated that chemotherapy-treated patients, irrespective of response, exhibited consistently low TIL levels (**Figure 1E**)(see Methods). Notably, chemotherapy responders included samples from both primary breast and metastatic lesions (stage IV), whereas chemotherapy non-responders were exclusively derived from primary breast tumors with stage II–III disease (**Supplementary Figure S1D**). When integrating peripheral immune profiling, functional subsets defined from PBMC decreased only in responders post-treatment in chemotherapy, consistent with systemic immune activation and potential redistribution toward tumor or other tissues (**Figure 1E**). In combination therapy, responders, predominantly from lymph node lesions (stage IV), also exhibited post-treatment decreases in circulating functional subsets, particularly in high TIL and PD-L1 tumors, further suggesting immune localization (**Figure 1D, E**)(**Supplementary Figure S1D**) (see Methods). In contrast, combination non-responders displayed post-treatment increases in peripheral immune subsets despite low TIL levels, indicating a disconnect between systemic activation and intratumoral recruitment, consistent with immune dysfunction or exclusion (**Figure 1E**)(**Supplementary Figure S1D**). Using linear mixed-effects modeling, we demonstrated that peripheral blood gene expression robustly predicted tissue-level expression (pre-treatment β = 0.49, t = 44.9; post-treatment β = 0.57, t = 54.6, p < 0.001), with significant interactions indicating that this relationship varies across immune functional subsets (**Supplementary Figure S1E**). These results support the hypothesis that systemic immune signatures reflect and are associated with tumor immune states in a subset-dependent manner. To further understand these dynamics, we examined changes within the lymphoid and myeloid subsets across treatment and response groups.

### 2.2. Lymphoid Compartments

The lymphoid compartment showed distinct composition and mechanistic patterns across treatments and responses. In chemotherapy responders, baseline priming of adaptive immunity was a key feature, whereas post-treatment persistence of effector/exhaustion states characterized resistance, suggesting these patients might benefit from immunotherapy. In combination therapy, baseline effector/exhaustion signatures were associated with response, while resistance was linked to dysregulated adaptive immunity.

#### 2.2.1. CD4 T cell subsets: baseline immune priming characterize responders

Of the six CD4^+^ T cell clusters identified (see Methods, **Supplementary Table S2**), four functional clusters (Activated naïve-like CD4, regulatory CD4 (T_reg_), T-helper 1 (Th1), and T stem-like memory (CD4_Tsm) that were representative across patients were selected for detailed analysis (**Figure 2A, B**). In the chemotherapy group, responders exhibited a baseline increase in activated naïve-like CD4 with a high naïve score, enriched for interferon-driven inflammatory program and antigen processing, features indicative of enhanced immune priming (**Figure 2C-F**) (6). Notably, this aligns with prior observations by Zhang *et al.* (7,8) where they identified a naïve CD4 cluster (CD4 Tn_CCR7) as predictive of response to both PTX and Nab-PTX. CD4_Tsm subsets showed elevated activation and anti-apoptotic scores, consistent with a primed and long-lived memory phenotype (**Figure 2C-F**). T_reg_ were also increased, exhibiting upregulated TCR signaling, stress response, adhesion, metabolism, and pro-apoptotic scores, indicative of their tumor reactivity before treatment (**Figure 2C, F**). In the combination therapy group, responders also exhibited baseline increase in activated naïve-like CD4 and CD4_Tsm, mirroring the immune priming signatures observed in the chemotherapy groups (**Figure 2C**).

**Figure 2.**
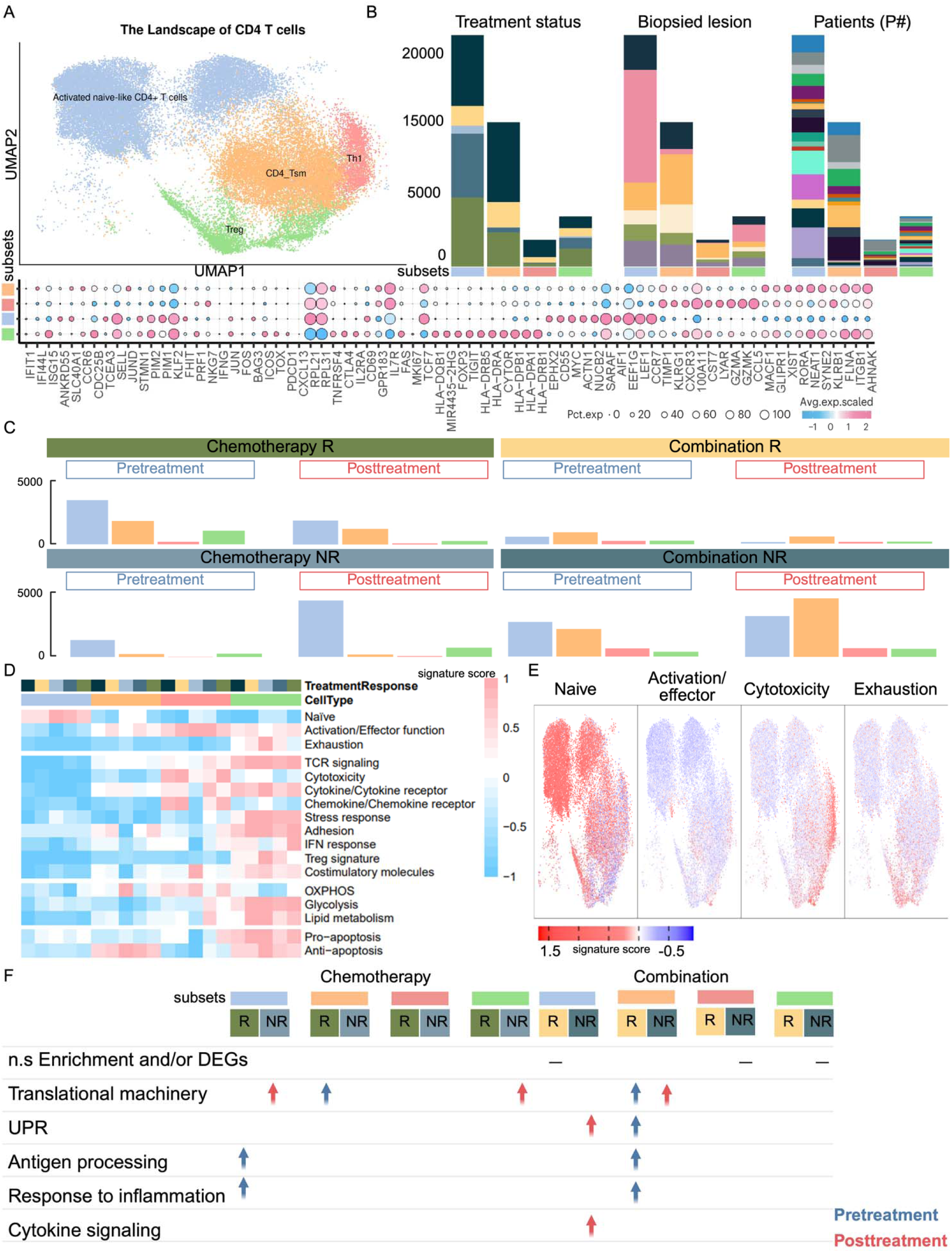
Comprehensive landscape of CD4D T cell subsets in blood samples and their distribution across conditions. **(A)** UMAP representation of CD4 T cell sub-clusters in blood, colored by distinct functional states (top). Bubble plot showing representative marker gene expression across clusters (bottom). Additional marker genes are listed in **Supplementary Table S2**. **(B)** Stacked bar plots of CD4 T cell sub-clusters across treatment and response groups, biopsied lesions, and individual patients, using color annotations consistent with **Figure 1B**. **(C)** Bar plots comparing the distribution of CD4 T cell sub-clusters before and after treatment across samples, colored by functional states. **(D)** Heatmap of 16 curated gene signature scores across CD4 T cell clusters (23) **(E)** Expression of four representative gene signatures selected from (**D)**. **(F)** Table showing enriched pathways or functional gene signatures associated with CD4 T cell sub-clusters, stratified by condition (blue: enriched in pre-treatment; red: enriched in post-treatment). This Figure highlights the transcriptional heterogeneity of CD4 T cells in blood, their patient-specific variability, and dynamic changes in sub-cluster composition and function in response to treatment. Pathway enrichment analyses point to key functional programs shaping peripheral CD4 T cell dynamics.

#### 2.2.2. CD4 T cell subsets: metabolic dysregulation characterize non-responders

In contrast with responders, there was a lack of increased immunity at baseline of non-responders. In chemotherapy non-responders, treatment induced expansion of activated naïve-like CD4 with high naïve scores, alongside Treg enriched for TCR signaling, stress response, adhesion, glycolysis, and Treg-associated scores, indicating a shift towards regulatory and pro-tumor activation (**Figure 2C-E**). Both CD4 subsets were also enriched for translational machinery, further supporting a regulatory and metabolically active immune state (**Figure 2F**). Notably, while our observations are from peripheral blood, they align with findings by Zhang et al. who reported increased T_reg_ frequencies post-chemotherapy in tumor tissue non-responders, suggesting a systemic and intratumoral response associated with treatment resistance (7,8). In combination therapy, non-responders, post-treatment CD4_Tsm expanded with high anti-apoptotic scores alongside enrichment for translational programs. Activated naïve-like CD4 exhibited high naïve scores (**Figure 2C-E**) and were enriched for cytokine signaling and activation of the unfolded protein response (UPR), indicative of cellular stress response rather than effective priming (**Figure 2F**). Increased frequencies of T_reg_ with elevated TCR signaling, cytokine receptor, adhesion, and apoptotic scores, as well as Th1 cells with activation, cytotoxicity, chemokine receptor, and OXPHOS scores, reflected an imbalanced and potentially dysfunctional CD4 T cell response (**Figure 2C-E**).

#### 2.2.3. CD8 T cell subsets: baseline naïve and effector expansion in chemotherapy and terminal differentiation in combination therapy characterize responders

Of the fifteen CD8^+^ T cell clusters identified (see Methods, **Supplementary Table S2**), seven functional clusters (CD8_naïve, CD8_naïve + CD8_naïve_248+, effector cells (CD8_eff), Activated CD8_eff, exhausted T cells (CD8_Tex), Proliferating CD8_eff + interferon stimulating genes (ISG), CD8_Tex-prog) that had cell contributions from all patients were selected for detailed analysis (**Figure 3A, B**). In the chemotherapy group, responders exhibited a baseline increase in several naïve clusters (CD8_naïve and CD8_naïve + CD8_naïve_248+) with increased naïve scores and enrichment in translational and antigen processing programs **(Figure 3C-G, Supplementary Table S2**). The effector CD8^+^ subsets, including CD8_eff, activated CD8_eff, proliferating CD8_eff + ISG, and CD8_Tex-prog, were elevated, highlighting the association of transitional and cytotoxic states with chemotherapy response (**Figure 3C**). CD8_eff cells showed elevated activation, chemokine receptor, and adhesion scores (**Figure 3D, E**). Differential analysis further confirmed upregulation of translational machinery, cell migration, cytoskeleton remodeling, and oxidative phosphorylation (OXPHOS) (**Figure 3G**). Activated CD8_eff were enriched for TCR signaling, cytokine and chemokine receptor, and MAPK signaling scores (**Figure 3D, E**). Proliferating CD8_eff + ISG subsets were marked by high stress, metabolic, and pro-apoptotic scores, and CD8_Tex-prog displayed increased exhaustion, TCR signaling, IFN response, and pro-apoptotic scores (**Figure 3D, E**). Expansion of effector and transitional CD8+ T cells with heightened TCR signaling and metabolic activity is consistent with antigen-experienced cytotoxic immune programs during chemotherapy, a pattern consistent with established models with metabolic reprogramming in activated CD8+ T cells (9). In combination therapy responders, CD8^+^ T cell dynamics showed distinct features compared to chemotherapy alone. Responders displayed increased frequencies of CD8_naïve with high naïve score, alongside elevated effector subsets (CD8_eff, activated CD8_eff, and CD8_Tex), indicating a functionally engaged and terminally differentiated cytotoxic program (**Figure 3C-E**). While CD8_eff cells largely mirrored the activation, TCR, cytokine/chemokine receptor, and adhesion patterns seen with chemotherapy, they also showed higher pro-apoptotic scores (**Figure 3D, E**). Differential analysis revealed similar enrichment in cell migration and cytoskeletal remodeling pathways, with combination therapy uniquely adding inflammatory response programs (**Figure 3G**). Notably, CD8_Tex cells in the combination setting exhibited elevated activation, exhaustion, MAPK, and adhesion scores (**Figure 3D, E**). These systemic profiles are consistent with previously reported intratumoral CD8 CXCL13 exhausted T cells (7), suggesting a coordinated peripheral and tissue immune engagement.

**Figure 3.**
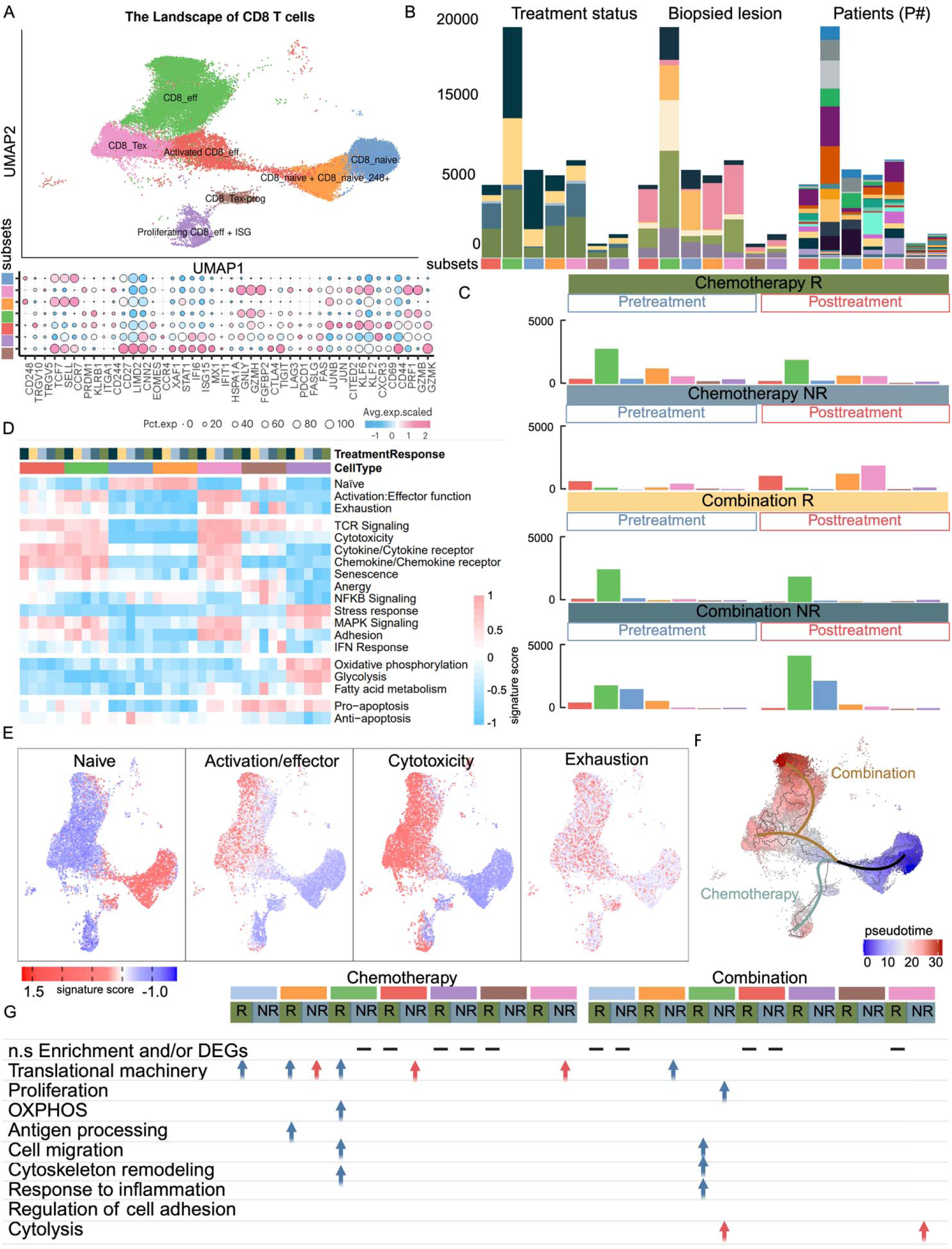
Comprehensive landscape of CD8D T cell subsets in blood samples and their distribution across conditions. **(A)** UMAP representation of CD8 T cell sub-clusters in blood, colored by distinct functional states (top). Bubble plot showing representative marker gene expression across clusters (bottom). Additional marker genes are listed in **Supplementary Table S2**. **(B)** Stacked bar plots of CD8 T cell sub-clusters across treatment and response groups, biopsied lesions, and individual patients, using color annotations consistent with **Figure 1B**. **(C)** Bar plots comparing the composition of CD8 T cell sub-clusters in blood before and after treatment across multiple samples, colored by distinct functional states. **(D)** Heat map illustrating the expression of 19 curated gene signatures across CD8 T cell clusters. **(E)** Expression of four representative gene signatures selected from **(D)**. **(F)** Monocle 3 trajectory analysis of CD8 T cell differentiation revealing three main divergent trajectories. Cells are color-coded by pseudo time. **(G)** Table representation of enriched pathways or functional signatures associated with CD8 T cell sub-clusters, stratified by condition. (Blue: enriched in pre-treatment. Red: enriched in post-treatment. This Figure illustrates the heterogeneity of CD8 T cells in peripheral blood, their inter-patient variability, and dynamic reprogramming in response to treatment. Enrichment analysis highlights context-specific pathways associated with CD8 T cell functional transitions.

#### 2.2.4. CD8 T cell subsets: terminal differentiation and dysregulated cytotoxic programs in non-responders

In chemotherapy non-responders, treatment induced expansion of CD8_naïve + CD8_naïve_248+, characterized by high naïve scores (**Figure 3C-E**) and enrichment in translational pathways (**Figure 3G**). Additionally, effector and terminal subsets including CD8_eff, activated CD8_eff, proliferating CD8_eff + ISG, and CD8_Tex were expanded, reflecting a skew toward cytotoxic and fully differentiated exhausted phenotypes (**Figure 3C**). These subsets showed strong enrichment for TCR signaling, activation, cytotoxicity, MAPK signaling, cytokine/chemokine receptors, adhesion, and pro-apoptotic scores (**Figure 3D, E**). Notably, CD8_eff + ISG and CD8_Tex subsets demonstrated enhanced stress response and metabolic signatures, consistent with terminal differentiation and immune exhaustion (**Figure 3D, E**). Differential analysis showed enrichment of translational machinery associated with Activated CD8_eff and CD8_Tex (**Figure 3G**). In the combination therapy group, non-responders exhibited a distinct pattern of immune remodeling compared to chemotherapy. At baseline, CD8_naïve + CD8_naïve_248+ subsets were elevated and exhibited high naïve and NFKB scores (**Figure 3C-E**) and enrichment in translational pathways (**Figure 3G**). Additionally, activated CD8_eff subsets showed high activation, TCR signaling, cytokine and chemokine receptors, senescence, and MAPK signaling scores, suggesting dampened effector function prior to treatment (**Figure 3 C-E**). Post-treatment increases in CD8_naïve cells with high naïve scores, CD8_eff subsets enriched for activation, cytotoxicity, cytokine and chemokine receptors, and adhesion scores, and CD8_Tex with T cell receptor (TCR) signaling, cytotoxicity, cytokine and chemokine receptors, MAPK signaling, and adhesion scores (**Figure 3C-E**). Differential analysis confirmed strong enrichment of cytolysis-associated genes in effector subsets (**Figure 3G**). These results indicated ineffective or dysregulated T cell activation in the combination non-responder group.

Pseudo time analysis of CD8 T cells revealed two main trajectories: (1) from naïve through activated effector states toward transitional populations (Tex-prog, CD8_eff+ISG), and (2) toward fully differentiated exhausted (Tex) and cytotoxic effector subsets (**Figure 3F**). In the chemotherapy group, cells were primarily enriched in the transitional branch, whereas combination therapy cells predominantly occupied the terminally differentiated branch, reflecting more complete effector maturation under combination therapy. Notably, the apparent differences between responders and non-responders within each treatment group were largely explained by temporal dynamics, with responders enriched in these states at pre-treatment and non-responders at post-treatment.

#### 2.2.5. B cells: translationally active and primed at baseline in responders

Of the twelve B cell clusters identified (see Methods**, Supplementary Table S2**), three functional clusters (CD19_B cells, CD20_B cells, memory B cells (CD27_Bmem)) that were representative across patients were selected for detailed analysis (**Figure 4A, B**). In the chemotherapy group, responders exhibited a baseline increase in B cell populations, including CD20^+^ B cells, CD19^+^_AP1, and CD27^+^ memory B cells, enriched for cytoplasmic translation (**Figure 4D**), suggesting increased activation and protein synthesis (10). Pre-treatment PPI network analysis of these populations revealed enrichment of ribosomal and translational regulators, stress proteins, and BCR signaling components (CD79A/B, PTPN6), indicating a transcriptionally and translationally engaged B cell state in chemotherapy responders (**Figure 4E**). Notably, while these results are derived from peripheral blood, they align with prior findings demonstrating that B cells within the tumor microenvironment were predictive of response to chemotherapy: decreasing in paclitaxel-treated responders but increasing in those treated with nab-paclitaxel plus atezolizumab (8). Together, these data suggest that B cell activation signatures in circulation and tissue are associated with treatment responsiveness. In combination therapy responder, baseline B cell populations displayed distinct dynamics compared to chemotherapy. CD20^+^ B cells were enriched for translational machinery, B cell activation, and antigen processing, while CD27^+^ memory B cells were enriched for cytoskeleton remodeling and antigen processing, and CD19^+^_AP1 were elevated at baseline but lacked significant mechanism enrichment, suggesting differential B cell dynamics compared to chemotherapy (**Figure 4D**).

**Figure 4.**
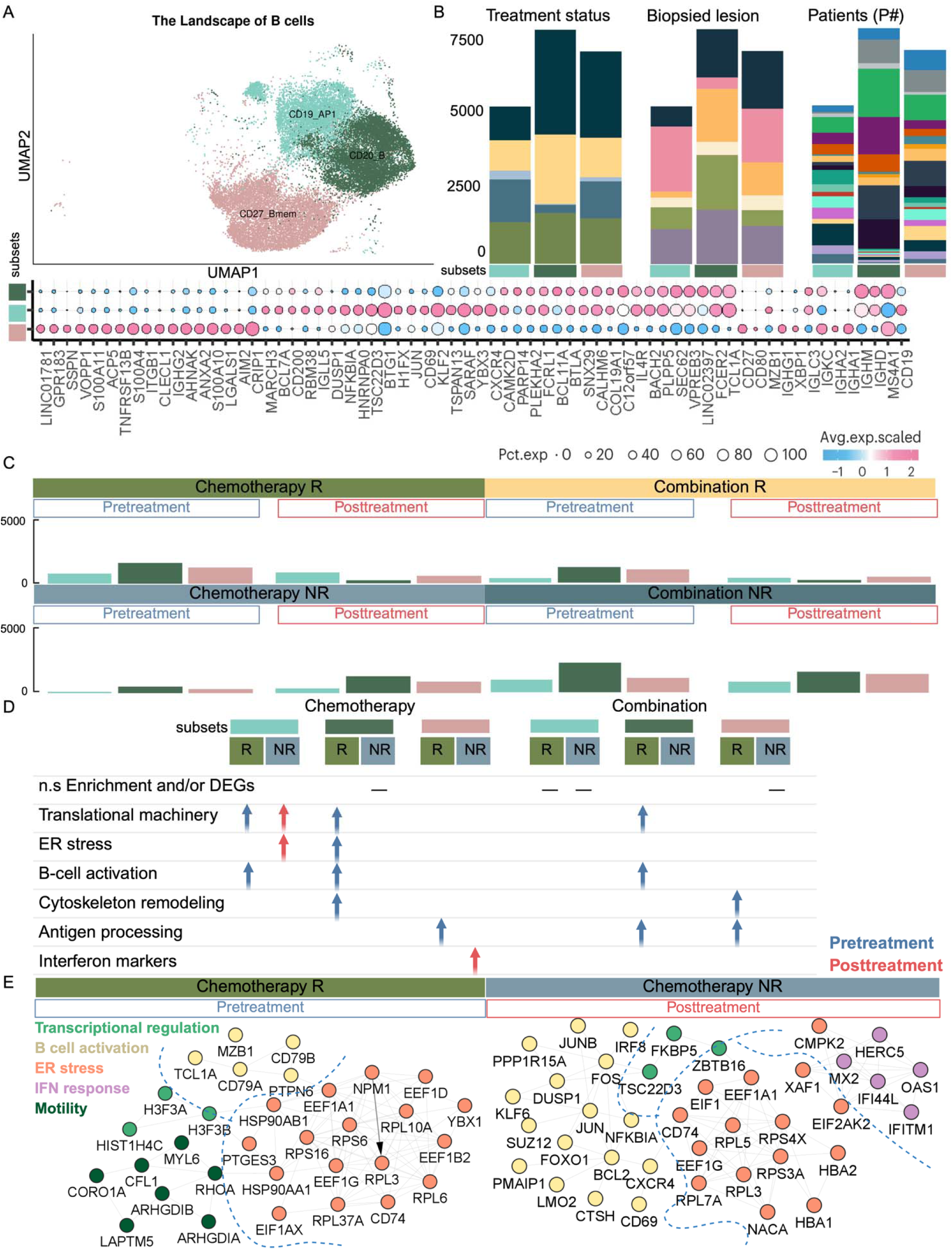
Comprehensive landscape of B cell subsets in blood samples and their distribution across conditions. **(A)** UMAP representation of B cell sub-clusters in blood, colored by distinct functional states. Th bubble plot shows representative marker gene expression across clusters. Additional marker genes are listed in **Supplementary Table S2**. **(B)** Stacked bar plots of B cell sub-clusters across treatment and response groups, biopsied lesions, and individual patients, using color annotations consistent with **Figure 1B**. **(C)** Bar plots comparing the composition of B cell sub-clusters in blood before and after treatment across multiple samples, colored by functional states. **(D)** Table summarizing enriched pathways or functional signatures associated with B cell sub-clusters, stratified by condition. Blue: enriched in pre-treatment; Red: enriched in post-treatment. **(E)** PPI networks highlight distinct mechanisms enriched in the Chemo-R pre-treatment (right) and Chemo-NR post-treatment (left) groups. This Figure illustrates the heterogeneity of B cells in peripheral blood, highlighting patient-specific variation and dynamic changes in sub-cluster composition and functional states in response to treatment. Enrichment and network analyses underscore key signaling pathways associated with B cell activity across clinical conditions.

#### 2.2.6. B cells: stress-regulated and dysregulated programs in non-responders

In chemotherapy non-responders, treatment induced expansion of B cell populations, including CD20^+^ B cells, CD19^+^_AP1, and CD27^+^ memory B cells, which were enriched for translational machinery (**Figure 4C, D**). PPI network analysis highlighted ribosomal and translational regulators, stress- and apoptosis-related proteins, and interferon-inducible genes. Immune checkpoint and regulatory-associated genes, including CD69, CXCR4, JUN, IRF8, and ZBTB16, were also notable, suggesting a multifaceted stress-regulatory B cell response (**Figure 4E**). In combination therapy non-responders, baseline frequencies of B cell subsets including CD20 cells enriched for translational programs, CD19 _AP1, and CD27 memory B cells were elevated. Among these, only CD19 _AP1 showed a higher frequency relative to post-treatment, indicating selective persistence of specific B cell populations (**Figure 4C, D**).

#### 2.2.7. Innate lymphoid cells: translational enrichment in responders versus exhausted phenotypes in non-responders

Of the thirteen NK cell clusters identified (see Methods**, Supplementary Table S2**), three functional clusters (NK1, NK3, NK8) that were representative across patients were selected for detailed analysis (**Figure 5A, B**). In chemotherapy responders, NK1 cells exhibited elevated translational activity and cytotoxic markers at baseline, suggesting an innate readiness that parallels CD8^+^ T cell activation (**Figure 5C, D**). This coordinated priming reflects classical NK cell function, part of the broader family of group 1 innate lymphoid cells (ILC1), which are known to provide rapid effector responses and support adaptive immunity (13). In contrast, combination therapy responders showed enhanced innate engagement following treatment, with expansion of NK1 cells with elevated translational and inflammatory signaling, indicative of heightened NK cell activation and functional differentiation (**Figure 5C, D**).

**Figure 5.**
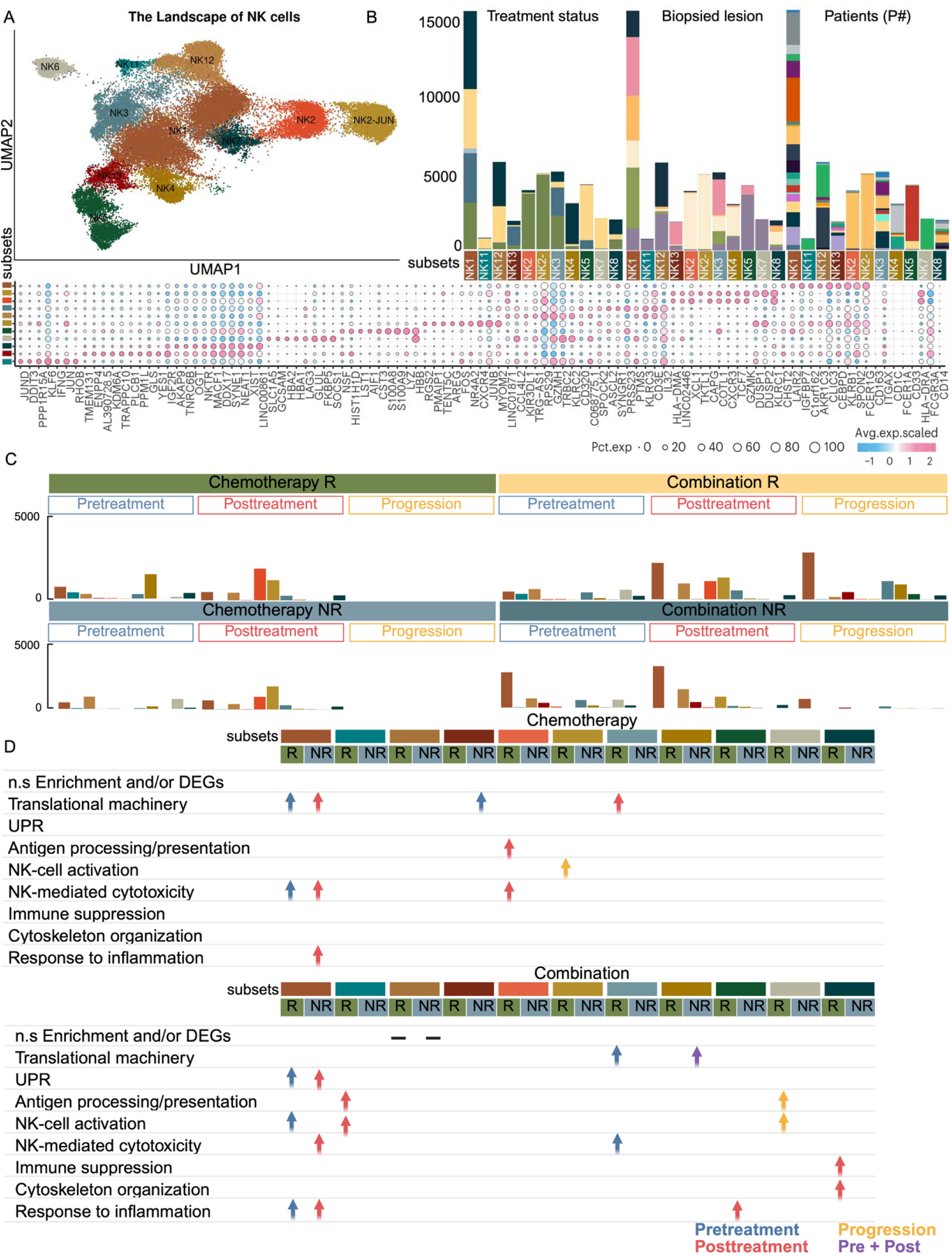
Landscape and dynamics of NK cell subsets in blood across treatment conditions. **(A)** UMAP of NK cell sub-clusters in blood, colored by functional states. Bubble plot show representative marker gene expression across clusters. Additional marker genes are listed in **Supplementary Table S2. (B)** Stacked bar plots showing NK sub-cluster distributions across treatment groups, response status, biopsied lesions, and individual patients, using color annotations consistent with **Figure 1B**. **(C)** Bar plots comparing NK sub-cluster compositions before and after treatment. **(D)** Table summarizing enriched pathways or functional programs by sub-cluster and condition. Blue: enriched in pre-treatment; Red: enriched in post-treatment; Yellow: enriched in progression; Purple: enriched in both pre- and post-treatment. This Figure highlights the heterogeneity of NK cells, their patient-specific patterns, and dynamic functional shifts in response to treatment. Enrichment analyses reveal key immune pathway underpinning NK cell responses in the blood.

In chemotherapy non-responders, treatment induced an increase in NK1 cells enriched for translational machinery, inflammatory response, and cytotoxicity/exhaustion markers, while NK3 subsets were enriched for translational activity (**Figure 5C, D**). Collectively, these patterns suggest that chemotherapy in non-responders promotes immune subset expansion, but skewed toward a dysregulated, pro-inflammatory, and exhausted phenotype lacking coordinated effector function (14). Similarly, in combination therapy non-responders, NK3 cells at baseline exhibited cytoplasmic translation enrichment and inhibitory markers (KIR2DL3, KLRG1), consistent with an exhausted phenotype (**Figure 5C, D**). Following treatment, NK1 cells showed increased ER stress, inflammatory response, and cytotoxic/exhaustion signatures, paralleling the chemotherapy non-responder profile (**Figure 5C-E**).

These findings highlight distinct patterns of immune dysregulation between treatment groups. In chemotherapy non-responders, CD8+ T cell diversity expanded but lacked coordinated functional engagement with an elevated exhaustion phenotype. In contrast, combination-treated non-responders exhibited more defined CD8+ activation but were marked by CD4+ stress responses, suggesting dysfunctional immunity. This aligns with observations of a pan-cancer CD4 T cell stress–response state linked to immunotherapy resistance (15). Notably, translationally active CD20+ and CD27+ B cell subsets remained elevated post-treatment, but lacked functional enrichment, suggesting limited contribution to effective immunity. Both treatments showed elevated exhaustion levels.

### 2.3. Myeloid cells

Myeloid responses showed treatment- and response-specific temporal immune programs. Chemotherapy responders displayed baseline enrichment of metabolically active, antigen-presenting myeloid cells, whereas combination responders exhibited post-treatment expansion of pro-inflammatory monocytes. Non-responders in both regimens showed post-treatment metabolic activation without effective inflammatory programs, paralleling dysfunctional lymphoid immune programs.

#### 2.3.1. Divergent Timing of Myeloid Immune Programs in Responders to Chemotherapy and Combination Therapy

Of the nine myeloid clusters identified (see Methods**, Supplementary Table S2**), three functional clusters (classical monocyte (CM), non-classical monocyte (NCM), dendritic cells (DC)) that were representative across patients were selected for detailed analysis (**Figure 6A, B**). In the chemotherapy group, baseline samples from responders showed increased frequencies of CMs enriched for translational machinery, and DCs enriched for translational machinery, ER stress, OXPHOS, and interferon signaling. This transcriptional profile is consistent with a metabolically active, antigen-presenting phenotype (**Figure 6C, D**). This suggests that pre-existing immune activation, particularly metabolically active and antigen-presenting myeloid cells, is necessary for an effective response to chemotherapy (6). Together with adaptive immune priming, this suggests an interplay between the adaptive and innate immunity, priming the microenvironment for tumor clearance. However, no significant expansion of myeloid cells was observed following chemotherapy treatment, similar to the lack of expansion observed in lymphoid cell types. In the combination therapy group, baseline samples from responders showed increased DC infiltration, with the transcriptional landscape showing enrichment for translational machinery, stress, and inflammatory response (**Figure 6C, D**). Following treatment, responders exhibited an expansion of CM enriched for chromatin remodeling, suggesting a memory-like phenotype with a pro-inflammatory transcriptional program (**Figure 6C, D**). NCMs were enriched for cytotoxic, interferon-responsive, and migratory phenotype, including ITGA4, TLN1, MSN, CORO1B markers (**Figure 6C, D**). These findings in parallel with reduced adaptive immunity are consistent with robust immune remodeling following anti-PD-L1 therapy.

**Figure 6.**
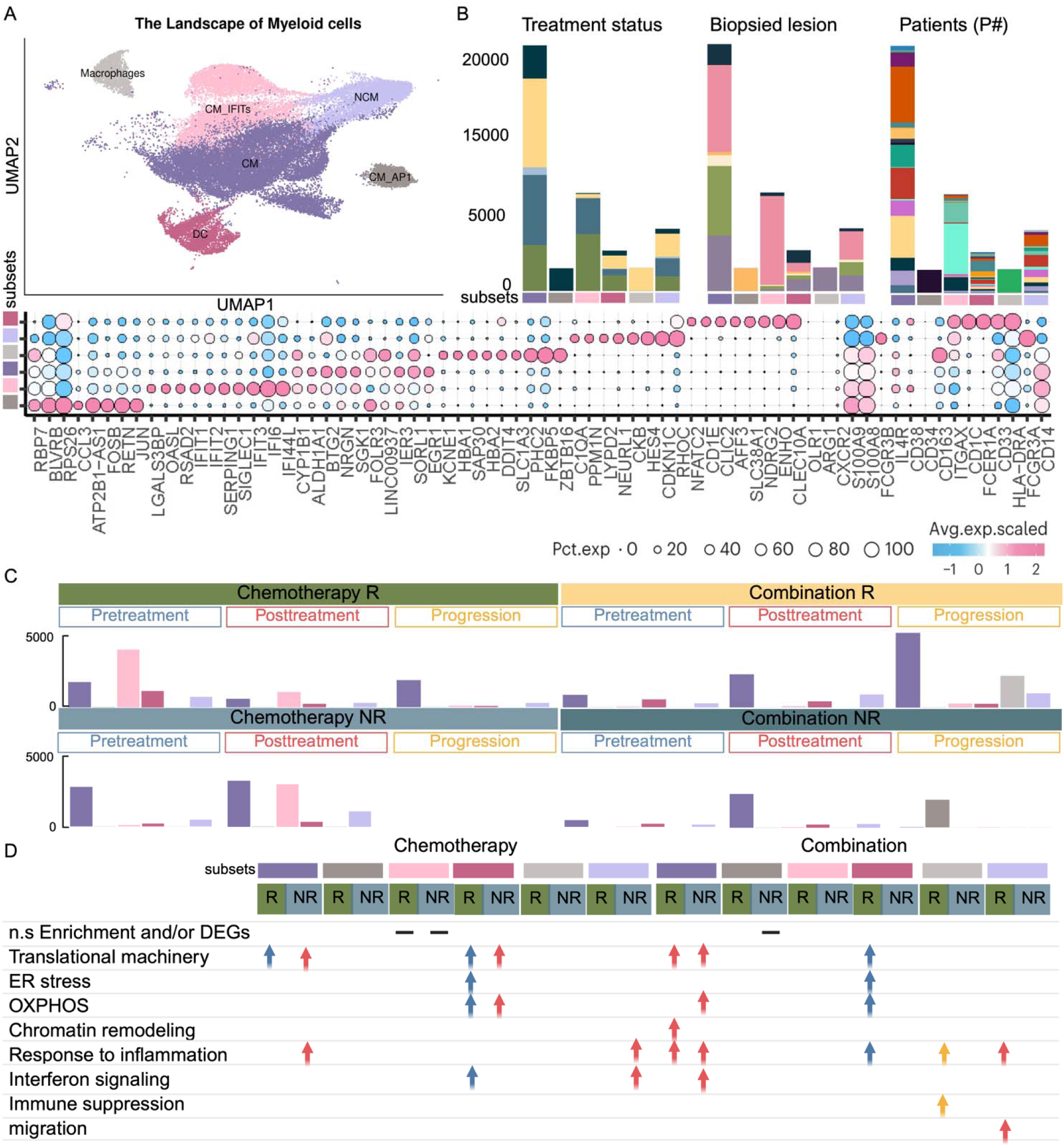
Landscape and dynamics of monocyte subsets in blood across treatment conditions. **(A)** UMAP of monocyte sub-clusters in blood, colored by distinct functional states. The bubble plot shows representative marker gene expression; additional markers are listed in **Supplementary Table S2**. **(B)** Stacked bar plots showing monocyte sub-cluster distributions across treatment groups, response status, biopsied lesions, and individual patients, using color annotations consistent with **Figure 1B**. **(C)** Bar plots comparing monocyte sub-cluster compositions before and after treatment. **(D)** Table summarizing enriched pathways or functional programs by sub-cluster and condition. Blue: pre-treatment; Red: post-treatment; Yellow: progression. This Figure highlights the heterogeneity of circulating monocytes, their patient-specific distribution patterns, and dynamic shifts in composition and function following treatment. Enrichment analyses reveal pathways underlying monocyte responses in the peripheral blood.

#### 2.3.2. Myeloid compartment in non-responders exhibits inflammatory skewing and immune dysfunction

At baseline, non-responder samples across both treatment groups showed a lack of higher myeloid presence than post-treatment. Following chemotherapy, non-responders exhibited an increased frequency of CMs enriched for cytoplasmic translation, inflammatory signaling, and wound-healing pathways (PF4, PF4V1, SPARC, TGFB1), indicative of a pro-tumorigenic and potentially immunosuppressive state. NCMs showed upregulation of inflammatory and type I IFN response programs, while DCs exhibited elevated cytoplasmic translation and OXPHOS, reflecting heightened metabolic activity (**Figure 6C, D**). Post-treatment, non-responders displayed a myeloid compartment dominated by metabolically active, pro-tumorigenic programs, in line with evidence that tumor metabolic reprogramming can induce suppressive myeloid phenotypes (16,17). Concurrently, the lymphoid compartment exhibited increased diversity, with features suggestive of T cell dysfunction/exhaustion. This pattern is consistent with delayed and ineffective immune activation, potentially reflecting tumor-driven immune dysregulation. A similar escape mechanism has been described in the context of tumor-mediated disruption of antigen presentation of T cell exhaustion as a mode of cancer immune evasion (18). In the combination group, post-treatment CMs upregulated cytoplasmic translation, OXPHOS, type I IFN signaling, and immunoregulatory genes (SOCS3, BCL3), suggesting activation skewed towards dysregulation rather than effective antitumor immunity (19) (**Figure 6C, D**). NCMs showed elevated cytoplasmic translation but lacked hallmark inflammatory or cytotoxic gene expression (**Figure 6C, D**), indicating a metabolically primed state without clear evidence of functional activation. Such metabolic reprogramming without effector function is a recognized pattern in tumor-associated myeloid cells, where translation-driven activation contributes to suppression rather than immune effector activity (20). In parallel, the lymphoid compartment exhibited an increase in terminally exhausted T cells, further reinforcing a state of ineffective immune activation in non-responders.

### 2.4. Tissue CD4+/CD8+ T cells dynamics diverge in non-responders

Of the seven CD4^+^ and fourteen CD8^+^ T cell clusters identified (see Methods**, Supplementary Table S2**), five CD4^+^ and four CD8^+^ functional clusters that were representative across patients were selected for detailed analysis (**Figure 7A, B, D, E**). At baseline, both chemotherapy and combination therapy responders showed enrichment of adaptive immune subsets, including CD4_naive, Tfh + Tex-prog, T_reg_, CD8_cytotoxic, CD8_Tex + MKI67, and CD8_naive-like + ISG populations, indicating a pre-existing, antigen-experienced T cell compartment poised for activation (**Figure 7C, F**). These observations alig with Zhang *et al.* (7,8), that identified subsets: naïve T cells, Tfh-CXCR5, and Tex-MKI67, enriched at baseline in PTX responders but appeared post-treatment in Nab-PTX responders. Following treatment, both groups showed increased activated naïve-like CD4+ T cells, while combination therapy showed increased activated CD8-eff and chemotherapy showed CD4-CM (**Figure 7C, F**). Note that Zhang *et al.* (7,8) showed that activated naïve-like CD4 was associated with combination post-treatment in PTX, while pre-treatment in Nab-PTX. In non-responders, baseline profiles diverged by treatment. Combination therapy non-responders showed enrichment of CD4_naïve, CD4_CM, Treg, activated CD8_eff, and CD8_cytotoxic subsets (**Figure 7C, F**). Chemotherapy non-responders instead exhibited broad T cell expansion post-treatment (**Figure 7C, F**), consistent with Zhang *et al.*’s (7,8) observation of post-chemotherapy T_reg_ increases.

**Figure 7.**
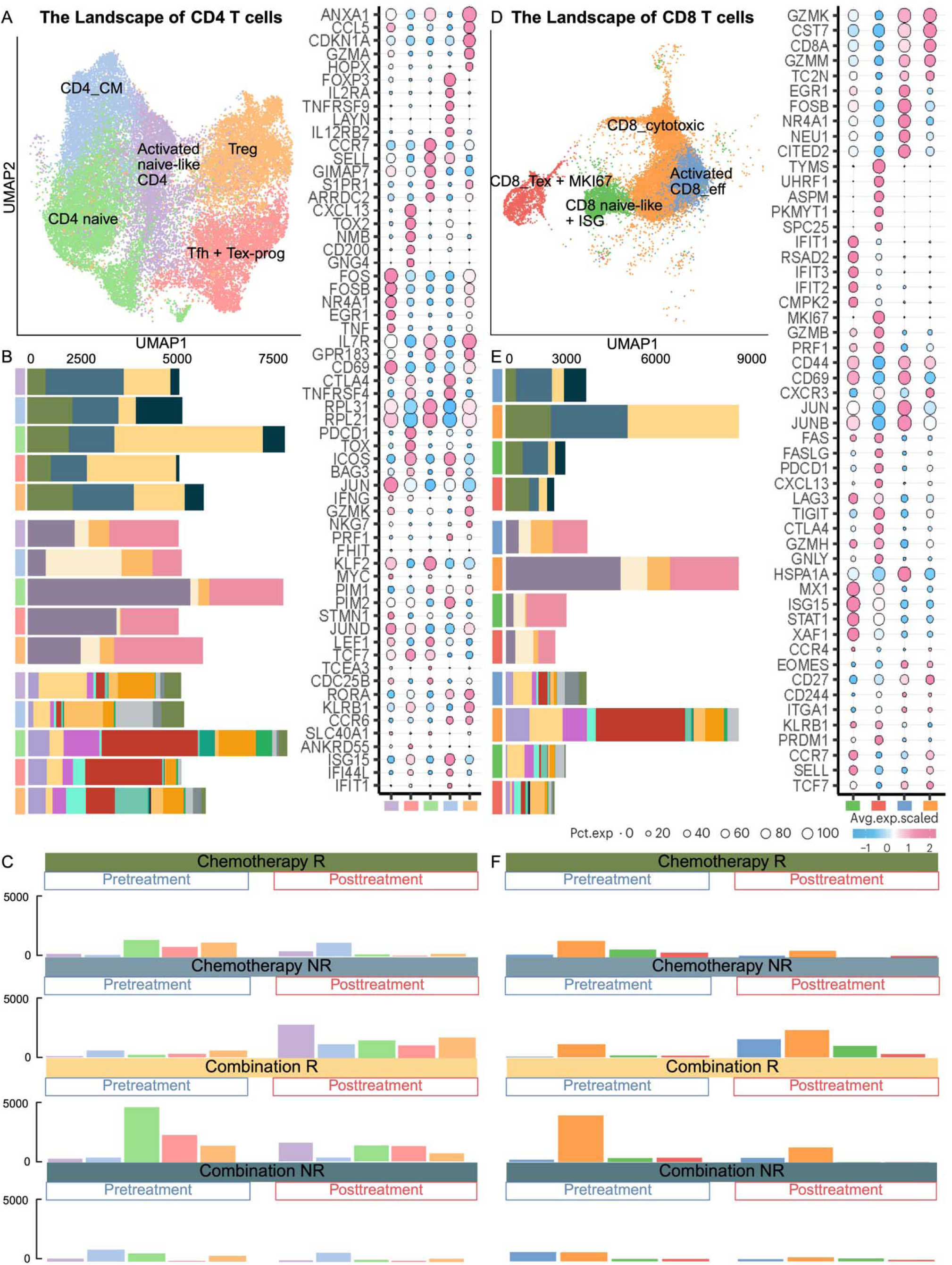
Comprehensive landscape of CD4+ and CD8+ T cell subsets in tissue samples and their distribution across conditions. **(A)-(D)** UMAP representation of CD4+ and CD8+ T cell sub-clusters in tissue, respectively, colored by distinct functional states. The bubble plot shows representative marker gene expression; additional markers are listed in **Supplementary Table S2**. **(B)-(E)** Stacked bar plots showing CD4+ and CD8+ sub-cluster distributions, respectively, across treatment groups, response status, biopsied lesions, and individual patients, using color annotations consistent with **Figure 1B. (C)-(F)** Bar plots comparing T cell sub-cluster compositions before and after treatment.

### 2.5. Coordinated systemic and tumor T cell programs distinguish chemotherapy and combination therapy responders

In chemotherapy responders, tissue PPI network analysis revealed enrichment of IFN-driven signaling, antigen processing, and strong exhaustion markers (**Figure 8A**). Blood profiling showed upregulation of T cell priming/activation states and early activation of CD8+ T cells (**Figure 9A**). This concordance suggests a sequential immune mobilization model, in which activated T cells detected in the blood are associated with intratumoral cytotoxic and exhaustion programs. In combination therapy responders, tissue networks highlighted ribosomal proteins, metabolic enzymes, and TCR activation markers, consistent with sustained antigen-driven effector function supported by high biosynthetic and metabolic capacity (**Figure 8B**). Baseline blood profiles mirrored this state, showing more fully differentiated and engaged adaptive immunity with active TCR signaling and cytotoxicity responses (**Figure 9B**), indicating parallel activation of systemic and intratumoral T cell compartments at treatment onset. Notably, in combination therapy, post-treatment blood shifted toward expansion of innate immune subsets with JAK–STAT, TNF, and IFN pathway activation. These observations are consistent with systemic innate immune re-engagement that may follow effective adaptive tumor clearance, even if not captured in matched tissue samples due to lack of representative innate immunity across patients (**Figure 9B**).

**Figure 8.**
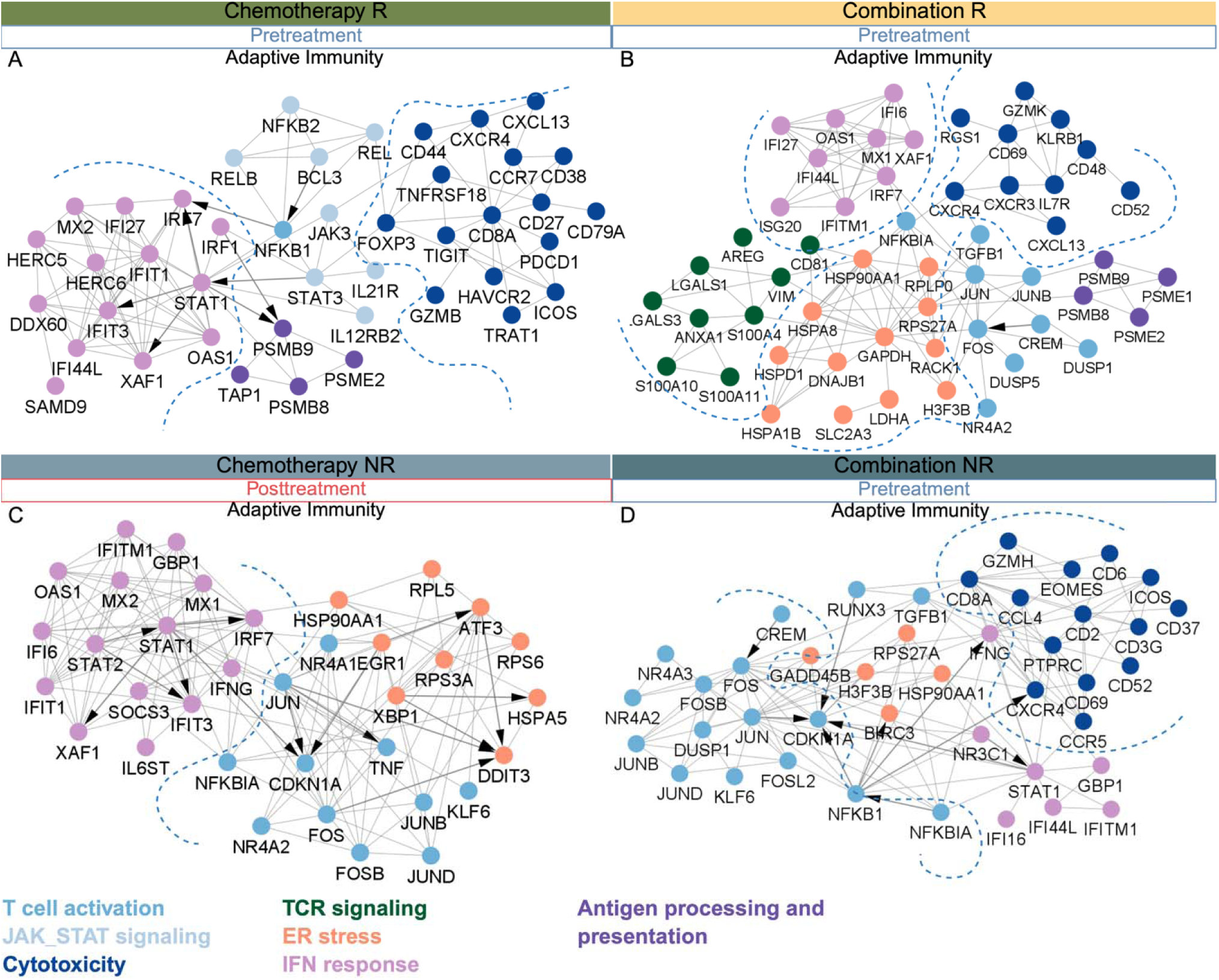
Protein–protein interaction (PPI) networks of treatment- and response-specific mechanisms in tissue. PPI networks were constructed from differentially expressed genes identified in each treatment and response group when comparing post- versus pre-treatment samples (see Methods). **(A)** Chemotherapy responders (pre-treatment): modules enriched in T cell activation, antigen processing, interferon signaling, and cytotoxicity. **(B)** Combination responders (pre-treatment): modules enriched in interferon signaling, T cell activation, ER stress, and TCR signaling. **(C)** Chemotherapy non-responders (post-treatment): modules enriched in T cell activation, ER stress, and interferon response. **(D)** Combination non-responders (pre-treatment): modules enriched in T cell activation, interferon signaling, ER stress, and cytotoxicity. Node color denotes functional modules (see legend) and arrows indicate transcription–target relationships. Together, thes networks reveal condition-specific immune regulatory circuits and highlight potential therapeuti targets.

**Figure 9.**
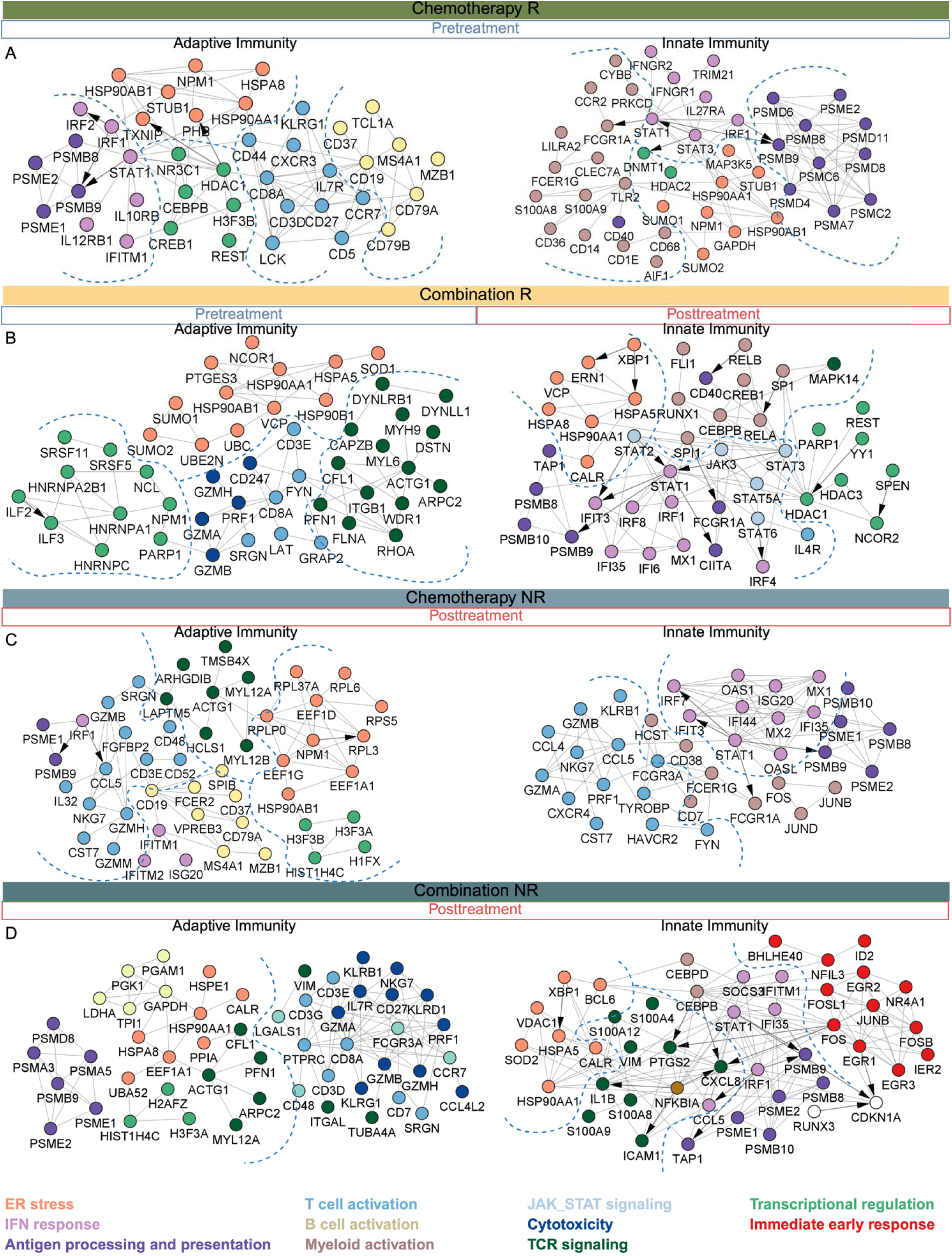
Protein–protein interaction (PPI) networks of treatment- and response-specific mechanisms in blood. PPI networks were constructed from differentially expressed genes identified in each treatment and response group when comparing post- versus pre-treatment samples (see Methods). **(A)** Chemotherapy responders (pre-treatment): adaptive immunity (left panel) modules enriched in T cell activation, antigen processing, interferon signaling, and transcriptional regulation. Innate immunity (right panel) modules enriched in myeloid activation, interferon signaling, ER stress and antigen processing **(B).** Combination responders (pre-treatment): adaptive (left panel) modules enriched in T cell activation, cytotoxicity, ER stress, and transcriptional regulation. Combination responders (post-treatment): innate immunity (right panel) modules enriched in myeloid activation, interferon signaling, ER stress, and transcriptional regulation. **(C)** Chemotherapy non-responders (post-treatment): adaptive immunity (right panel) modules enriched in T- and B-cell activation, TCR signaling, ER stress, and interferon response. Innate immunity (right panel), myeloid activation, JAK-STAT signaling, antigen processing, and interferon signaling. **(D)** Combination non-responders (post-treatment): adaptive immunity (left panel) modules enriched in T cell activation, interferon signaling, ER stress, and cytotoxicity. Innate immunity (right panel) modules enriched in motility, antigen presentation, interferon signaling, ER stress and immediate early response. Node color denotes functional modules (see legend) and arrows indicate transcription–target relationships. Together, these networks reveal condition-specific immune regulatory circuits and highlight potential therapeutic targets.

In chemotherapy non-responders, post-treatment showed tissue enrichment for antigen-driven activation and effector differentiation, but also stress-response and immunoregulatory programs (TNF, TNFSF10, IL6ST, NFKBIA, SOCS3, ZC3HAV1) (**Figure 8C**). Blood analysis revealed parallel induction of effector, cytotoxicity, and exhaustion markers, reinforcing the interpretation that transcriptionally primed T cells remain functionally constrained (**Figure 9C**). In tissue, combination therapy non-responders at baseline displayed PPI networks combining antigen-driven activation, cytotoxic function, and metabolic adaptation, but these were accompanied by immunoregulatory and stress-response signatures (TNFAIP3, BIRC3, TGFB1, NR3C1, NFKBIA) and chemokine/cytokine modulators (CCR5, CXCR4, CCL4) (**Figure 8D**), suggesting functional suppression within an immunosuppressive microenvironment (21). This suppression was mirrored in blood, where reduced effector signatures were observed (**Figure 3**). Across both therapies, non-responders demonstrated post-treatment expansion of innate immune subsets in blood, characterized by oxidative stress, metabolic dysfunction, and immunosuppressive signaling. PPI analysis indicated upregulation of antioxidant and inflammatory regulators (SOD2, PRDX, HMOX1) alongside NFKBIA and PTGS2, with stronger induction in combination non-responders, suggesting unresolved inflammation and dysfunctional innate immune re-engagement (**Figure 9C, D**).

### 2.6. Patient-specific immune dynamics

While group-level analyses revealed broad immune differences between responders and non-responders, patient-specific analyses uncovered nuanced immune adaptations shaped by metastatic site and therapeutic response. In chemotherapy responders, patients P020 (breast), P022 (chest wall metastasis), and P013 (liver metastasis) exhibited distinct myeloid and NK cell dynamics (**Figure 5B**) (**Figure 6B**). In blood, a baseline increase in CM_IFIT cells was consistently observed across these patients, reflecting a shared inflammatory signature (**Figure 6C**). Interestingly, in P013, treatment was followed by an increase of NK2 and at progression, increased by NK2_JUN, both involved in processing and presenting antigens, suggesting a dynamic NK cell response (**Figure 5C-E**). Tissue profiling of P013 revealed a sequential immune trajectory. Baseline pro-inflammatory monocytes transitioned into pro-inflammatory macrophages post-treatment, followed by increased NK1, NK2, TAM + ISG, macrophages, and macrophage + LYVE1 (**Supplementary Figure S2A, C**) (**Supplementary Figure S3A, C**). These populations displayed elevated expression of inflammatory and cytokine response genes, emphasizing the role of innate immunity in mediating chemotherapy response in metastatic disease (22) (**Supplementary Figure S2D**) (**Supplementary Figure S3D**). Notably, baseline adaptive immune engagement, characterized by primed CD4+ T cells and diverse CD8+ effector subsets, preceded the expansion of innate immune populations, suggesting a coordinated interplay in which early adaptive activation primes subsequent innate immune responses during treatment and disease progression.

In chemotherapy non-responders, blood samples from patients P018, P023, P024, and P025 (all breast) showed a post-treatment increase in CM_IFIT, aligning with the inflammatory profile observed at the group level (**Figure 6C**). In combination therapy responders, patients P019 and P007 (LN metastases) exhibited distinct immune dynamics. In blood, both patients showed post-treatment increases in NK5 and NK11, subsets associated with NK activation and antigen processing, consistent with group-level observations and highlighting innate immune activation following combination therapy (**Figure 6C, D**). This aligns with group-level findings, reinforcing the role of innate lymphoid immune activation following combination therapy. At progression, NK6, NK12, and macrophage populations were elevated, indicating ongoing immune adaptation (**Figure 6C**). In contrast, combination therapy non-responders, including P005 and P017 (both with liver metastases), showed distinct baseline immune features. In PBMC analysis, NK4 emerged in a patient-specific manner, present at baseline in P005 and post-treatment in P017, suggesting differential NK cell adaptation (**Figure 5B, C**). Tissue profiling of P017 revealed baseline enrichment of NK5, characterized by cytoplasmic translation and cytoskeleton organization (**Supplementary Figure S2B, C**), alongside CMs characterized with early immune activation, which were replaced post-treatment by macrophage + HS3ST2 expressing immunosuppressive markers (**Supplementary Figure S3B, C**). P005 showed NK3 cells enriched for cytoplasmic translation at baseline (**Supplementary Figure S2B, C**). Additionally, P016 (chest wall metastasis) showed increased NK12 at baseline, with NK7 expressing cytoskeleton organization and checkpoint markers (**Supplementary Figure S2B, C**), and emergence of CM_AP1 emerging at progression in PBMC (**Figure 6B**). Tissue samples revealed early macrophage + CCL3 enrichment, suggesting a pre-existing myeloid-driven immune phenotype (**Supplementary Figure S3B, C**). Collectively, these findings suggest that innate immune subsets with immunosuppressive features are associated with poor response to combination therapy.

Together, these findings highlight the heterogeneity of immune responses across patients and therapeutic regimens. While adaptive immunity appears to play a critical role in initiating response, the dynamic engagement or suppression of innate immune subsets during treatment and progression may be key to sustaining or modulating therapeutic efficacy. These results underscore the importance of incorporating patient-specific immune profiling to refine stratification strategies and inform the development of individualized therapeutic approaches.

### 2.7 Extension to I-SPY2 clinical trial data

To contextualize these findings against publicly available datasets, we leveraged scores derived from the I-SPY2 clinical trial (11). Correlation analysis demonstrated that NK1, Treg, and activated/exhausted CD8 programs positively aligned with tumor chemokine, interferon, and T/B-cell modules, while proliferating ISG-CD8 programs showed consistent negative association, indicating concordance between systemic immune states and established tumor immune phenotypes (**Supplementary Figure S4A**). To maintain treatment consistency, analyses were restricted to the pretreatment paclitaxel arm. Univariate logistic regression showed directional but underpowered associations with pCR. In contrast, multivariable LASSO modeling identified a coordinated peripheral immune signature associated with response, negative contributions from Activated naïve-like CD4 and dendritic programs, and positive contributions from NK, naïve CD8, and Treg states, suggesting composite systemic immune readiness may distinguish responders (**Supplementary Figure S4B**). Permutation testing indicated that the multivariate model performed better than random expectation but did not reach statistical significance (p=0.128), consistent with limited cohort size and correlated immune features.

## 3. Methods

### 3.1. Data overview

Published scRNA-seq data from Zhang et al. (GEO accession: GSE169246) which included 489,490 cells from 22 advanced TNBC patients, spanning 75 tissue (primary/metastatic) and blood samples were downloaded. The samples were collected at baseline, 4 weeks post-treatment, and at progression (**Supplementary Table S1**). Patient P002 was excluded as indicated in Zhang et al (7). The final dataset used for all downstream analyses included tissue samples (n = 15) and blood samples (n = 20). Patients had received either paclitaxel alone (chemotherapy; tissue n = 7, blood n = 10) or paclitaxel plus atezolizumab (combination therapy; tissue n = 8, blood n = 10), with documented treatment response available for most cases. Clinical response, including responder and non-responder, tumor-infiltrating lymphocytes (TIL), and PD-L1 scores, were downloaded from the original study.

### 3.2. Data filtering, scaling and dimensionality reduction

Data was filtered by excluding cells based on the following criteria: more than 10% mitochondria read, less than 500 expressed genes, less than 500 detected transcripts, and a complexity (log10 genes/UMI) of less than 80%. Following this QC, 432,837 cells remained for downstream analysis. Tissue-derived and PBMC samples were processed independently using Seurat (v.5.3.0), resulting in two data objects comprising 143,085 cells (tissue) and 289,752 cells (PBMC) across 27085 features/genes. Data scaling and dimensionality reduction was performed per vignette, with the optimal dimensions for clustering identified using *ElbowPlot* (tissue nPCs = 30 and PBMC nPCs= 31). The cells were clustered using the Louvain algorithm and visualized using UMAP.

### 3.3. Detecting cell types (Cluster annotation)

Cell type identities for each cluster detected was established using a threefold strategy: (1) SingleR with multiple reference datasets (DatabaseImmuneCellExpressionData, MonacoImmuneData, NovershternHeamtopoieticData), (2) differential expression testing via Seurat’s *FindAllMarkers* (Wilcoxon, logFC > 0.25, min.pct > 0.25, Bonferroni < 0.05), and (3) canonical marker expression and legacy knowledge. Markers were visualized using feature and bubble plots (top DEGs), enabling lineage assignment and transcriptional state assessment. Clustering yielded distinct immune populations in tissue samples, including 15 T cell clusters, 5 B cell clusters, 2 NK cell clusters, and 8 myeloid clusters. PBMC samples revealed 13 T cell clusters, 4 B cell clusters, 5 NK cell clusters, and 9 myeloid clusters.

### 3.4. Profiling T cell subsets

From the dataset, CD4 and CD8 T cells were extracted from both tissue and PBMC compartments for higher-resolution analysis. In tissue, a total of 90,286 CD4 /CD8 T cells (comprising 3 initial CD4 clusters and 12 initial CD8 clusters) were subsampled and re-clustered into 7 CD4 and 14 CD8 transcriptionally and functionally distinct subsets. In PBMC, 120,213 CD4 /CD8 T cells (from 9 initial CD4 clusters and 14 initial CD8 clusters) were subsampled and re-clustered into 6 CD4 and 15 CD8 distinct subsets. Subsampling was performed to ensure balanced representation of patient samples and avoid bias from uneven cell recovery. Cell type annotation was guided by established lineage and functional markers, including canonical naïve, central memory, effector, cytotoxic, proliferative, and regulatory T cell signatures (**Supplementary Table S2**) (1,2,23,24). To assess the transcriptional characteristics of various T cell subsets, we evaluated curated functional scores (e.g., exhaustion score, cytotoxicity score, anergy score), which were established based on a predefined set of genes obtained from a published pan-cancer T cell study (23), using gene module scoring (*AddModuleScore* function) in Seurat. Each score was calculated as the average expression level of the respective gene set.

### 3.5. Profiling B cell subsets

33,880 B cells (4 original clusters) were subset from the original data object, re-clustered, and grouped into 10 functionally distinct B cell states (plus 2 undefined), based on markers and pathway enrichment related to BCR signaling, MHC-II expression, IFN response, AP-1 transcriptional regulation, and metabolic/stress programs (**Supplementary Table S2**) (1).

### 3.6. Profiling NK cell subsets

In tissue, 9,332 NK cells (2 initial clusters) were subset from the original dataset and re-clustered into 8 transcriptionally and functionally distinct subsets based on DEG analysis and established classifications (1,2,23). In PBMC, 55,745 NK cells (5 initial clusters) were subset and re-clustered into 13 functional subsets using the same approach. Annotation incorporated canonical cytotoxic markers, activation and exhaustion profiles, and inflammatory gene signatures (**Supplementary Table S2**).

### 3.7. Profiling Myeloid subsets

In tissue, 17,866 myeloid cells (8 initial clusters) were subset and re-clustered into 23 transcriptionally distinct subsets. In PBMC, 52,949 myeloid cells (9 initial clusters) were subset and re-clustered into 9 subsets. Annotation identified classical (CD14) and non-classical (CD16) monocyte populations, with further subdivision guided by transcriptional heterogeneity and functional gene programs, including inflammatory signaling, stress responses, and metabolic pathways (**Supplementary Table S2**).

### 3.8. Trajectory inference of CD8^+^ T cells

To reconstruct differentiation trajectories of CD8 T cells, we used Monocle 3 (v.1.3.7) (25–29). After dimensionality reduction with UMAP and clustering, we constructed cell trajectories by learning principal graphs using the *learn_graph* function with use_partition = FALSE to preserve global connectivity across all subsets. Default parameters were used, as these yielded biologically coherent trajectories, with naïve T cells at the origin and expected branching toward effector and exhausted states. Pseudotime was assigned using *order_cells*, anchoring the root node based on naïve marker expression.

### 3.9. Functional enrichment and PPI/TF network construction

Functional enrichment was identified via functional annotation clustering available through *clusterProfiler* (30) or *Enrichr* (31) in R. Gene Ontology (biological process and molecular function) pathway analyses was performed using *clusterProfiler’s EnrichGO* function. A network of high confidence (>0.70) human protein-protein interaction (PPI) was downloaded from STRING database v12 (32), containing 15,588 proteins and 236,930 interactions. Additionally, transcription factor (TF)-target data downloaded from TRRUST-db (33) was mapped onto the PPI interaction network to generate a custom PPI/TF network.

Therapy-, response, and timepoint-specific PPI/TF subnetworks were generated using DEGs from adaptive and innate immune subsets described earlier.

In blood these included:

- Adaptive Chemo Responder (Pre-treatment): 746 proteins, 8,108 interactions
- Innate Chemo Responder (Pre-treatment): 986 proteins, 6,876 interactions
- Adaptive Chemo Non-responder (Post-treatment): 176 proteins, 4,078 interactions
- Innate Chemo Non-responder (Post-treatment): 232 proteins, 3,560 interactions
- Adaptive Combo Responder (Pre-treatment): 361 proteins, 3,560 interactions
- Innate Combo Responder (Post-treatment): 2,089 proteins, 13,919 interactions
- Adaptive Combo Non-responder (Post-treatment): 103 proteins, 1178 interactions
- Innate Combo Non-responder (Post-treatment): 731 proteins, 5,101 interactions

In tissue, these included:

- Adaptive Chemo Responder (Pre-treatment): 211 proteins, 1,139 interactions
- Adaptive Chemo Responder (Post-treatment): 41 proteins, 274 interactions
- Adaptive Chemo Non-responder (Post-treatment): 239 proteins, 3,280 interactions
- Adaptive Combo Responder (Pre-treatment): 166 proteins, 2,579 interactions
- Adaptive Combo Responder (Post-treatment): 38 proteins, 101interactions
- Adaptive Combo Non-responder (Pre-treatment): 162 proteins, 2066 interactions Each network was refined by selecting DEGs, identifying associated TFs, and extracting their 1-step PPI neighbors. All networks were visualized, annotated, and analyzed using Cytoscape (34).

### 3.10. Statistical modeling of blood-tissue expression relationships

We used linear mixed-effects models (LMMs) to quantify the relationship between blood and tissue gene expression across immune subsets. Models were fitted in R using lme4 package. For each gene, expression in tissue was modeled as a function of blood expression, immune cell subsets, and their interaction, with a random intercept for the patient. Models were fit separately for pre- and post-treatment data using restricted maximum likelihood (REML). Model performance was summarized by marginal and conditional R^2^ values using the performance package. To assess robustness, we performed leave-one-patient-out analyses by iteratively removing one patient, refitting the model, and recording (i) changes in REML criterion values, (ii) shifts in fixed-effect estimates, and (iii) variance components. The fixed-effect regression coefficient (β) quantifies the expected change in tissue gene expression per one-unit increase in blood gene expression, adjusted for immune cell subset composition.

### 3.11. External validation using I-SPY2 peripheral transcriptomic data

ISPY2 clinical trial raw data was downloaded from GSE194040, which includes pretreatment tumor gene expression profiles and clinical annotations for patients enrolled across multiple neoadjuvant treatment arms (11). Analyses were restricted to the paclitaxel arm to ensure treatment comparability with the discovery cohort. Gene expression data generated on Agilent microarray platforms (GPL20078) were downloaded from GEO. Probe-level measurements were mapped to gene symbols using platform annotations and collapsed to gene-level expression using median aggregation. Genes with zero variance across samples were excluded. Peripheral immune subset gene signatures derived from PBMC profiling in this study were projected onto the I-SPY2 cohort using single-sample gene set enrichment analysis (ssGSEA) implemented via the GSVA framework, generating per-sample immune program scores. Associations between immune subset scores and pathological complete response (pCR) were evaluated using logistic regression. Multivariate relationships were assessed using LASSO-regularized logistic regression with cross-validation, and model robustness was evaluated using permutation testing.

## 4. Discussion

This study provides a systematic characterization of immune cell populations and their associated mechanisms, revealing shared and distinct immunological features underlying treatment response and resistance in advanced TNBC patients receiving chemotherapy or chemo-immunotherapy. Peripheral immune dynamics differed by treatment and response, revealing therapy-specific immunological states with predictive and mechanistic interpretability. Chemotherapy responders had higher baseline immune cell counts than combination therapy responders, followed by post-treatment depletion consistent with known cytotoxic effects (41, 42). In non-responders, combination therapy patients showed higher baseline counts than chemo-only, but immune levels remained elevated post-treatment across groups. These findings are concordant with recent I-SPY2 peripheral blood reporting that therapeutic response is linked to systemic immune activation and cytotoxic remodeling in chemo-immunotherapy(12).

Across both treatment groups, our results highlight the importance of the dynamic coordination between adaptive and innate immune programs in modulating treatment outcomes. Enrichment of pathways including cytoplasmic translation, stress response, inflammatory signaling, cytoskeletal remodeling and migration, and metabolic programs including OXPHOS were observed in both responders and non-responders, underscoring their association with treatment outcomes. Post-transcriptional mechanisms, including the translation of mRNA, are critical contributors to rapid adaptive changes in the cellular proteome during immune responses(35). For instance, cytosolic protein translation is essential to increase glucose metabolism and degranulation capacity upon TCR activation, regulating the effector functions of CTLs and influencing the formation of memory cells (42, 43). Additionally, reprogramming cellular protein synthesis in innate immunity plays a pivotal role in inflammatory responses to pathogens and tumorigenesis by enhancing DNA-dependent cGAS activation, which drives the expression of ISGs (36).

Chronic antigen exposure can induce PD-L1 expression in tumor cells (likely via IFNG secreted from T cells) (37), and sustain PD-1 signaling in T cells, leading to an epigenetically imprinted exhaustion program (38–40). The presence of more differentiated T cells in combination therapy responders reflects this exhaustion phenotype and suggests higher tumor PD-L1 levels, potentially enhancing sensitivity to anti-PD-L1 therapy. This was supported by our observation that in responders tumors in combination were PD-L1^+^, while in non-responders they were PD-L1^-^. In contrast, chemotherapy responders showed less differentiated and more plastic T cells at baseline, consistent with lower PD-L1 expression and increased chemotherapy responsiveness. Combination therapy induced increased innate immune engagement, potentially in response to increased antigenic debris from tumor cell death. Such innate activation is critical for sustaining T cell memory and effector function, linking antigen presentation and cytokine signaling to adaptive immune maintenance (41). Innate immune reactivation post combination therapy may serve as a surrogate marker of response. Overall, these results provides a system-level context for previous studies showing that baseline immune cell diversity plays a crucial role in treatment outcomes (42–44).

In contrast, non-responders exhibited a myeloid compartment skewed toward an immunoregulatory, metabolically reprogrammed state, paralleled by increased dysfunctional/exhausted lymphoid populations across both treatments. Although canonical MDSC markers such as ARG1 were not prominently expressed, IL1B was present among DEGs, and coupled with upregulation of stress-adaptive and metabolic genes, this profile is consistent with an early-stage or atypical MDSC-like phenotype (45). This immunoregulatory shift may be associated with CD4 T cell polarization toward suppressive (T_reg_) or inflammatory (Th17) fates, as reported for metabolically reprogrammed human myeloid subsets (46). Such metabolically primed myeloid cells have been shown to suppress T- and NK cell activity or promote chronic inflammation, consistent with prior studies implicating monocyte and stromal metabolic reprogramming via cGAS–STING as a driver of therapy resistance (47). These findings highlight the plasticity of tumor-associated myeloid cells and the emerging importance of immunometabolism in shaping pro-tumoral functions (17,45).

Dysregulated NK cell responses emerged as a converging axis of resistance, with increased expression of DUSP1, CD81, and cytotoxic stress markers. Prior studies in TNBC have linked persistent NK activation with poor prognosis, with inflammatory NK signatures being increasingly recognized as pro-metastatic in solid tumors (46,48–50). Together, these data support a model in which adaptive-innate immune interplay in the peripheral blood is a marker and correlate of therapeutic outcome. Responders characterized by coordinated activation of lymphoid and metabolically competent myeloid populations, whereas non-responders display myeloid immunoregulation and lymphoid dysfunction or exhaustion. These results further highlight the potential of peripheral immune profiling to guide response prediction, resistance monitoring, and immunomodulatory strategies in TNBC.

In conclusion, our single-cell profiling of paired blood and tumor samples reveals that systemic and intratumoral immune remodeling distinguishes therapeutic responders from non-responders in TNBC. Responders exhibit coordinated activation of adaptive and innate immune programs, whereas non-responders display metabolic reprogramming and immune dysfunction across compartments. The implications of our study include potential development of blood-based biomarkers associated with treatment response, informing patient-specific therapeutic strategies, and informing the rational design of more precise combination therapies to increase response to treatment in TNBC patients.

## Acknowledgements

The authors thank Dr. Zemin Zhang and his group for their valuable guidance and support.

## Authors’ contributions

Z.M. analyzed the data, developed the models, wrote the first version of the manuscript along with preparing Tables and Figures. K.M. provided help with the design of the analysis and building the models. S.S. designed and supervised the project and revised the manuscript. All authors have read, edited, and accepted the manuscript.

## Data availability

All datasets analyzed in this study are publicly available in the Gene Expression Omnibus (GEO) under accession number GSE169246. Processed data and analysis scripts are available from the corresponding author upon request.

